# A Multimodal Neuroimaging Dataset to Study Spatiotemporal Dynamics of Visual Processing in Humans

**DOI:** 10.1101/2022.05.12.491595

**Authors:** Fatemeh Ebrahiminia, Morteza Mahdiani, Seyed-Mahdi Khaligh-Razavi

## Abstract

We describe structural and multimodal functional neuroimaging data collected from 21 healthy volunteers. Functional magnetic resonance images (fMRI) and electroencephalography (EEG) signals were acquired in separate sessions from the same individuals while they were performing a visual one-back repetition task. During functional sessions, participants were presented with images from five categories, including animals, chairs, faces, fruits, and vehicles. The stimulus set and experimental parameters were chosen to be similar to that of an available electrocorticography (ECoG) dataset, therefore creating a unique opportunity to study vision in humans with multiple complementary neuroimaging modalities. Individual-specific head models can be constructed for each participant using T1-weighted MPRAGE images and the recorded positions of the EEG electrodes. By combining the three functional modalities and the structural data, this dataset provides a unique setting to explore spatiotemporal dynamics of invariant object recognition in humans. This multimodal data can also be used to develop new methods for combining fMRI and electrophysiological modalities to come up with more accurate spatiotemporally resolved maps of brain function, which is inaccessible by any of the modalities alone.

## 1. Background & Summary

We recorded event-related fMRI and EEG data separately with the same stimulus set and similar experimental design used to record the ECoG data as described in Liu et al. (2009)^1^. Participants were presented with visual stimuli and were engaged in a one-back repetition task. The stimulus set included images from five categories: animals, chairs, faces, fruits, and vehicles. There were five different identities in each of the five categories, with five different sizes and orientations (=125 stimuli). Structural MRI and digitized electrode locations are also included in this dataset to construct subject-specific head models.

EEG and fMRI are complementary non-invasive neuroimaging techniques for studying human brain activity. FMRI data benefit from a higher spatial resolution, compared to EEG. In fMRI, we have access to the localized brain activity, allowing for regional-based analysis. However, in fMRI –being an HRF (hemodynamic response function)-based measurement – the detected signals are the result of the convolution of neural activity with the brain vascular response. This leads to a poor temporal resolution, therefore missing the underlying brain temporal dynamics^2^. EEG, on the other hand, has a rich temporal resolution, and is sensitive to the electrochemical current flows within and between the brain cells, detecting subtle electrical activities on the skull. Unfortunately, this comes at the cost of a poor spatial resolution^3^.

Current brain imaging techniques have been short in developing a noninvasive technique with whole brain coverage that can capture spatiotemporal dynamics of human brain activity with high resolution. However, several attempts have been made to develop algorithms for combining available neuroimaging techniques with high temporal (e.g. EEG or MEG) or high spatial resolution (e.g. fMRI) to investigate spatiotemporal dynamics of brain function^4–7^. When fusing these modalities, it is pivotal to investigate whether the combination of these techniques provides a more realistic view of the underlying brain activity. A promising approach is to compare the results obtained by M/EEG-fMRI fusion with the spatiotemporal neural dynamics obtained by invasive cell recordings, such as ECoG. In ECoG electrophysiological signals are recorded through intracranial electrodes implanted on the cortex. This technique is a powerful tool to separate neural events by their location and study fine task-related temporal dynamics, albeit with the caveat of not having the whole brain coverage^8^.

The presented dataset has the potential to be used for studying category selectivity and invariant object recognition with both high spatial and temporal resolutions, combining data from three modalities: EEG, fMRI and ECoG. This is a unique combination of modalities to study the human visual system. In addition, one may explore the validity of the existing EEG source localization techniques^9^ by comparing their outcome to the corresponding brain activity map obtained by fMRI and/or ECoG data. The shared task between the EEG, fMRI and ECoG data gives the opportunity to develop new EEG-fMRI fusion methods and test the outcome by drawing a comparison with the corresponding ECoG data.

## 2. Methods

### 2.1 Participants

The neuroimaging data were collected from a population of 21 healthy volunteers, 13 females and 8 males with an age range of 19 to 30 years (mean = 24.61). Participants were all university students, who responded to an invitation announcement to take part in the study. For all subjects, we collected one session fMRI and one session EEG data separately.

The study was conducted in accordance with the Declaration of Helsinki, and was approved by the ethics committee in Iran University of Medical Sciences. All participants got a health checkup before each session and signed a written consent form to participate in the study and also declared their consent for their anonymized data to be publicly available.

### 2.2 Stimulus set

The stimulus set for both EEG and fMRI included 25 objects from five categories (i.e. animals, chairs, faces, fruits, and vehicles), each shown at one of the following five transformations: (1) 3° visual angle, default rotation (i.e. 0°), (2) 1.5° visual angle, default rotation, (3) 6° visual angle, default rotation, (4) 3° visual angle, 45° rotation in depth, (5) 3° visual angle, 90° rotation in depth. This is the exact same stimulus set that was used to record the ECoG data in Liu et al. (2009)^1^. A subset of images is shown in Fig. S1. B (Liu et al, 2009)^1^. We used MATLAB 2016 ^10^ and the Psychophysics Toolbox ^11–13^ to code the experiment and present the stimuli to the subjects.

### 2.3 EEG experimental design and data acquisition

The EEG experiment was held in a room with mid lighting. A monitor was placed in front of the subject. The height of the subject’s chair was set so that the virtual line connecting the center of the monitor to the subject’s forehead had a slope of zero.

There were 32 runs. The images were presented once in each run in a random order. Each image was presented for 200 ms followed by an interstimulus interval (ISI) of 600 ms overlaid on a gray background. Participants were asked to try not to move their head through the runs. A fixation cross was used to help the subjects to fixate on the center of the screen. Each run took about 7 minutes, and the subjects were allowed to take rest between every four runs. To catch participants’ full attention, they were asked to perform a one-back repetition task, indicating whether the presented exemplar was repeated regardless of scale or viewpoint changes. They pressed a predefined key on a keyboard with their right index finger.

An 80-channel g. HIamp amplifier and g.GAMMAsys cap with g.LADYbird electrodes were used to record 64-channels EEG signals. The continuous EEG was digitized at 1100 Hz without applying any online filters. The signals were recorded with reference to the left mastoid. A ground electrode was placed at the forehead (Fpz), and three electrodes were used to record vertical and horizontal eye movements. Onset of trials and subject’s responses were marked by sending triggers via a parallel port. We used the Localite TMS navigator v3.0.33 with a Polaris Spectra camera to record the location of each electrode by placing the pointer on the scalp in the middle of each electrode.

### 2.4 FMRI experimental design and data acquisition

SIEMENS MAGNETOM Prisma 3 Tesla with a 64-channel head coil was used to collect the MRI data. T1-weighted MRI images were recorded using the MPRAGE sequence. Structural MRI parameters were: voxel size = 0.8×0.8×0.8 mm, slices per slab = 208, TR = 1800 ms, TE = 2.41ms, flip angle = 8°. The third dimension of DICOM files (i.e. slices per slab) for all subjects is 208, except for subject #2 (192 slices), subject #3 (192 slices), subject #20 (207 slices), and subject #21 (209 slices).

The functional MRI data were acquired using the following EPI sequence: voxel size = 3.5×3.5×3.5 mm, TR = 2000 ms, TE = 30 ms, flip angle = 90°, FOV read = 250 mm, FOV phase = 100%. Thirty four (34) interleaved slices were obtained in the order of even then odd slices. The whole experiment included 8 runs and took about 63 minutes. 237 volumetric images were acquired in each run. There were 125 image trials and 32 null trials in each run (each trial lasting 3 seconds). Each image was presented for 200 ms in a random order, followed by a blank screen for the remaining duration of the trial. Participants were given a response box to press a key with their right index finger, performing a one-back repetition task, indicating whether an exemplar was repeated regardless of scale or viewpoint changes. MRI head cushions and pillows were used to comfort participants and minimize head movements. To dampen the scanner noise, all subjects were provided with a set of earplugs. The stimulus set was presented to the subject through a mirror placed in the head coil.

## 3. Data Records

The EEG and fMRI data are available in Brain Imaging Data Structure^14,15^(BIDS) format on the Openneuro repository. Figure.1 shows the structure of the dataset directories and subdirectories stored on the repository. Our dataset consists of 21 downloadable folders, which are located in the main directory along with one *changes* file, one *README* file, a *dataset_description .json* file, a *participants .json* file, a *participant .tsv* file, another .*json* file that describes our task, and three other directories including *stimuli*, *derivatives*, and *sourcedata*.

**Figure 1.**
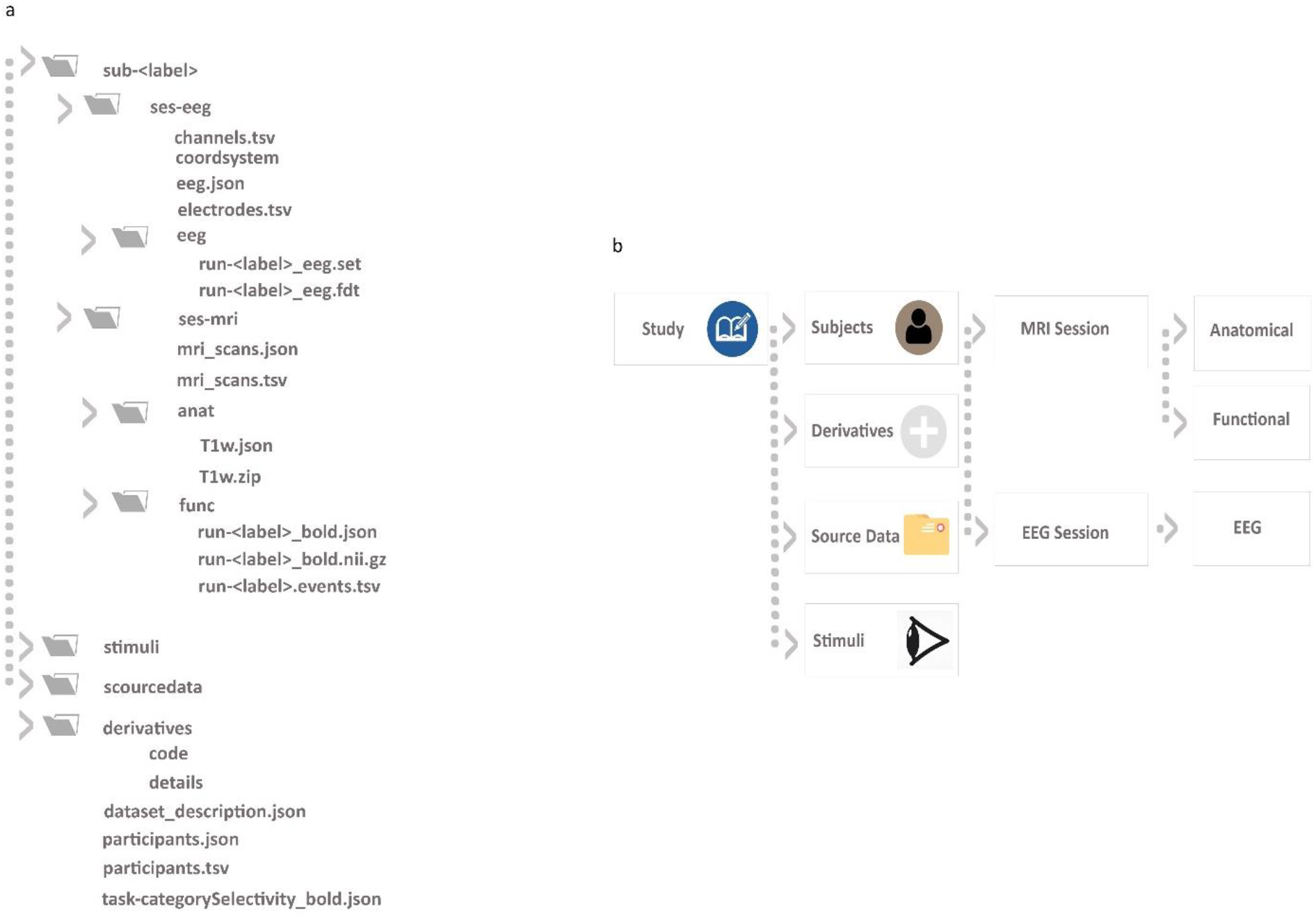
This figure shows the general view of the elements of the EEG-fMRI dataset in BIDS format. Main folders and subfolders along with their relevant metadata can be seen in this map. a) The structure of the uploaded files in BIDS format. Each directory that contains sub-directories or files is identified by a folder icon in its left side. b) The main subdirectory structure of the dataset is demonstrated in this section. Main folders that include the data and other files are arranged in 4 levels.

Each subject directory has two additional subdirectories, one for the EEG session and one for the fMRI session. The *ses-eeg* subdirectory includes a folder named *eeg* that has eight .*set* and eight .*fdt* files containing the EEG runs. In addition, there are *electrodes .tsv* files for the subject-specific location of EEG electrodes, *channels .tsv* file that includes information about EEG channels and two *. json* files: one for the coordinate system and one for the EEG metadata.

Another subdirectory in the subject folders is *ses-mri* that contains a .*tsv* and a .*json* file for metadata along with two directories: (1) *anat* directory that contains the anatomical data of the subject in the *nii.gz* format with its metadata file in *.json* format. (2) *func* directory that contains the EPI files for the eight runs in *nii.gz* format with their metadata as .*json* files and eight other *events .tsv* files. The *events .tsv* files include the exact timing of all the stimuli presented during each run.

As mentioned above, there are three additional directories in the parent folder: *stimuli, derivatives*, and *sourcedata*. In the *stimuli* directory, we have .*mat* files of the images presented to the subjects in the visual task. They are named according to the object’s category, identity, size, and orientation. The number after the category’s name identifies the object number. There are five different variations for each object identity. The codes *r1, r2*, and *r3* stand for the presentation size of 3° visual angle, with the following three rotations, respectively: default (i.e. 0°), 45°, and 90°. The *r4* and *r5* denote the presentation of stimuli in their default rotation (i.e. 0°), with the 6° and 1.5° visual angle respectively. For example, *face1r1ext_st.mat* refers to the first face identity from the face category, with the size of 3° visual angle and the default rotation (i.e. 0° rotation). The *derivatives* directory contains two subdirectories: *code* and the *details* folders. The code that was used to generate the *nii.gz* and .*json* files exists in the *code* folder. The *details* folder has the *info* subfolder including the folders specified with the number of subjects and has DICOM information of each subject. In addition to the *info* folder, the commands we used to produce fMRI BIDS as a text file are added to the *details* folder.

The *sourcedata* directory includes raw files from the mri scanner. The raw EEG data are not included as they are accessible in the *EEG file* for each subject. Inside *sourcedata* directory, there are 21 folders with the *sub-##* name format. Each subject folder includes functional and structural MRI DICOM data located in the *scans* folder. Table 1 summarize the demographic and the experimental information for the dataset.

**Table 1.** Participants’ demographic and the experiment’s information for EEG, fMRI, and ECoG dataset.

| Data type | Number of participants | Task | Gender | Age-range | Stimulus set | Source |
| --- | --- | --- | --- | --- | --- | --- |
| EEG | 21 | One-back repetition | 13 female (72%) | 18 to 30 | 125 images in 5 categories | Link (openfMRI) |
| fMRI | 21 | One-back repetition | 13 female (72%) | 18 to 30 | 125 images in 5 categories | Link (openfMRI) |
| ECoG | 11 | One-back repetition | 5 female (54%) | 12 to 34 | 125 images in 5 categories | External link |

\* 5 categories included animals, chairs, faces, fruits, and vehicles

## 4. Technical Validation

### 4.1 FMRI analysis

#### Univariate Analysis

We used MATLAB 2016 ^10^ and SPM12^16^ to preprocess the data. At first, we performed the slice timing correction (with the middle slice as the reference), and then the volumes from all runs were realigned with the first volume of the first run. To normalize the data to the standard MNI space, the structural images were aligned with the International Consortium of Brain Mapping (ICBM) space template with a voxel size of 2×2×2 mm, and the mean image over volumes was registered to the structural image, and the resulting transformations were applied to all realigned images. For univariate analysis, the functional data were spatially smoothed by a Gaussian kernel with a FWHM of 8 mm.

Time series of each voxel were concatenated over 8 runs, and the General Linear Model (GLM) was carried out on voxels’ time series for each subject. To this end, regressors were constructed by convolving a vector containing the onset of the stimulus with the canonical hemodynamic response function. To correct for residual movements, another 6 regressors per run were added. These regressors were the motion parameters: 3 rotations and 3 translations.

Figure. 2 shows the results of group level univariate analysis of all stimuli vs. null condition (t-test, p<0.001, uncorrected).

**Figure 2.**
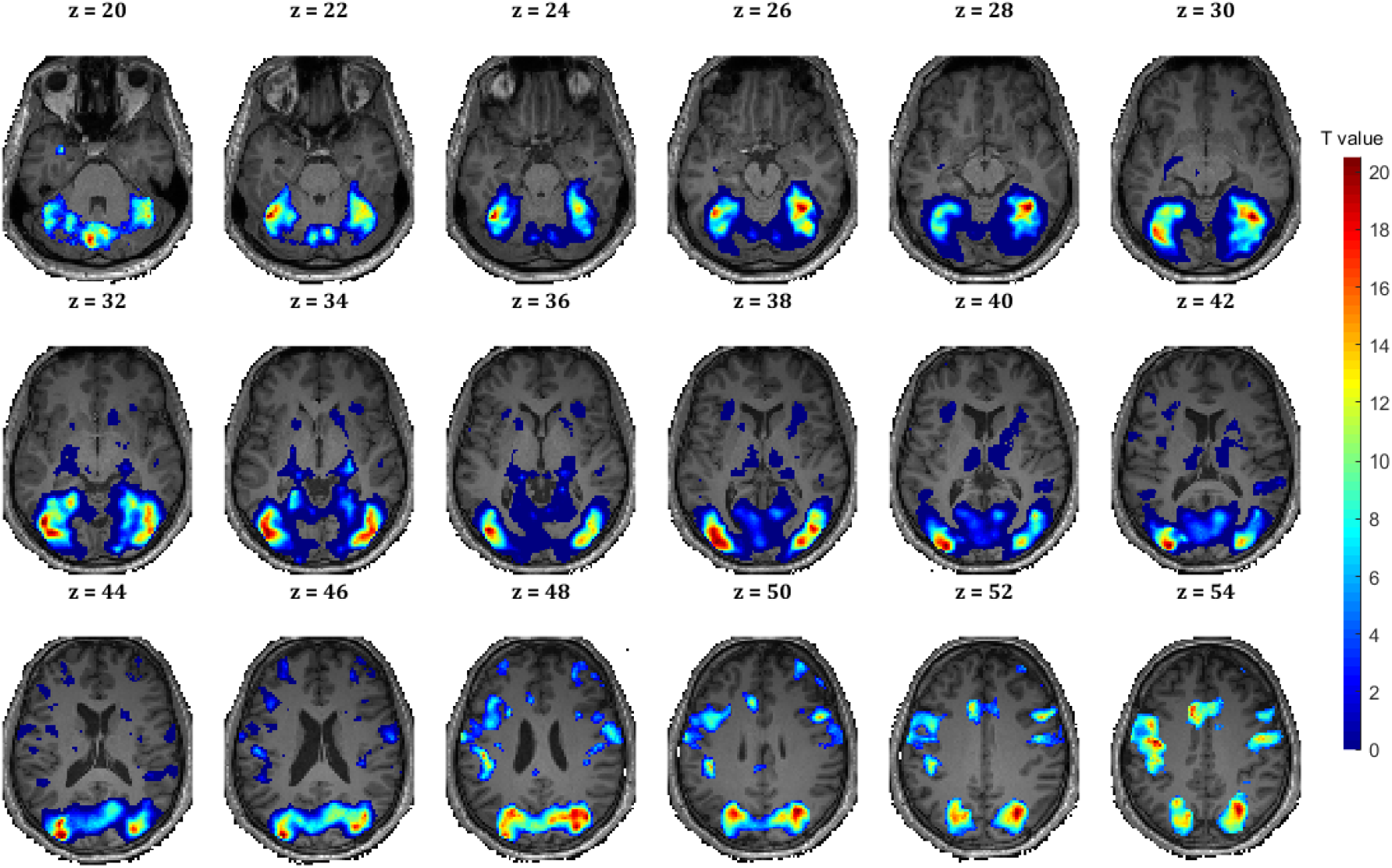
Group level univariate analysis of 21 subjects. Axial slices of activated voxels (all vs. baseline) are shown on a T1 image in the MNI space (p<0.001 uncorrected).

#### Multivariate Analysis

Multivariate pattern analysis with a searchlight approach^17^ was used to examine category selectivity. For every voxel in the brain, we extracted condition-specific voxel patterns in its vicinity (80 voxels) and used a SVM classifier to calculate the average pairwise decoding accuracy in each searchlight window for different categories (animals, chairs, faces, fruits, and vehicles). This procedure is applied to each subject’s beta-values obtained from the GLM analysis. For multivariate analyses, we used CoSMoMVPA^18^ and LIBSVM^19^ toolboxes in MATLAB 2016. The grand average performance for pairwise category selectivity is calculated by averaging accuracy maps for each subject and then averaging the results across all subjects.

To test the significance of the results, we used nonparametric statistics, right-sided signrank test, and corrected for multiple comparisons by FDR (p<0.05, across voxels) (Figure. 3)

**Figure 3.**
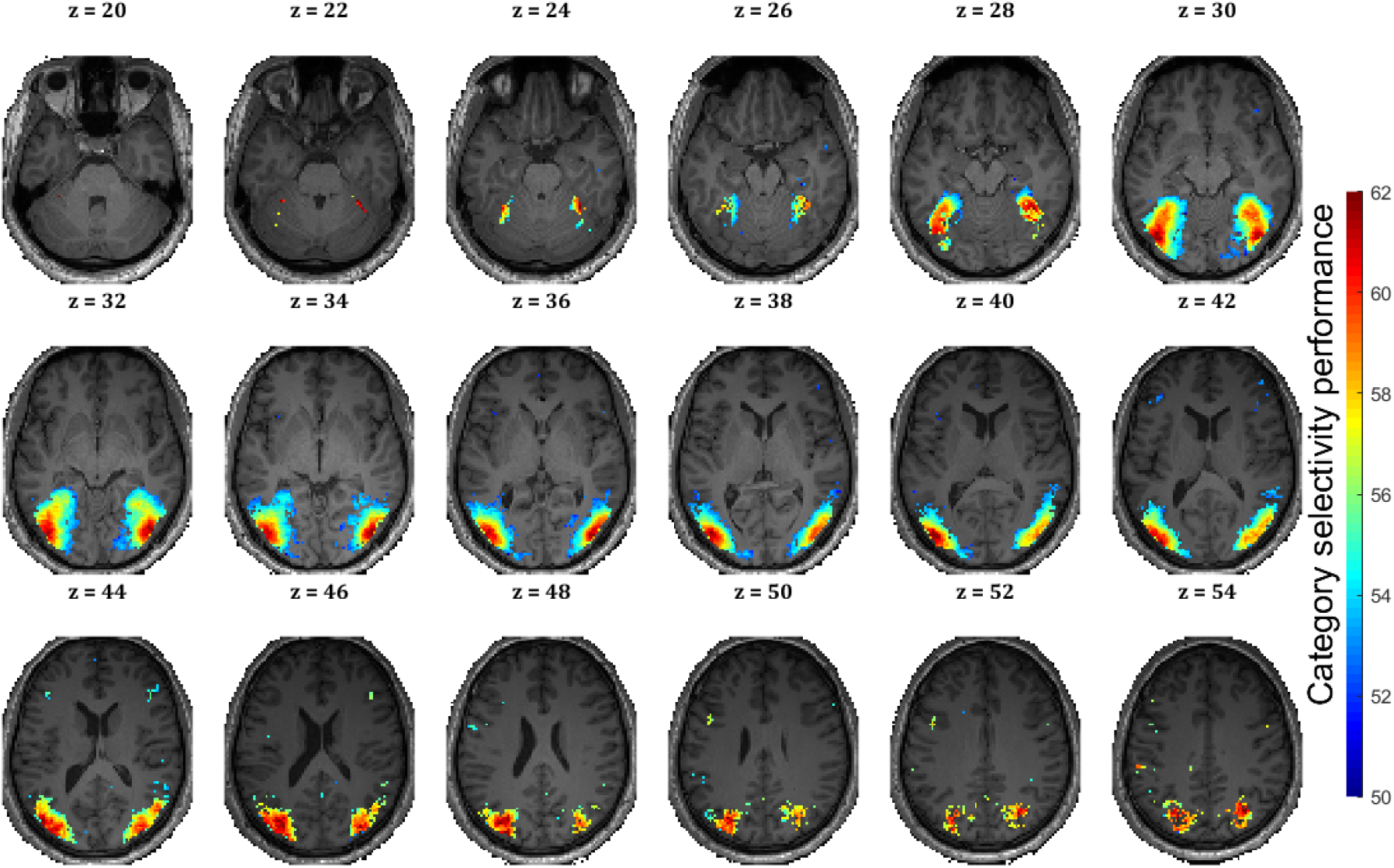
Multivariate pattern analysis of 21 subjects’ fMRI data. Regions with significant pairwise decoding accuracy (average across 21 subjects) are shown on a T1 image in the MNI space (one-sided sigrank test, FDR corrected at P < 0.05)

### 4.2 EEG analysis

#### Univariate Analysis

To analyze EEG data, we used EEGLAB^20^ in MATLAB 2016. The data from all runs were concatenated. The continuous data was filtered using a low-pass filter with a cutoff frequency of 40 Hz. We then resampled the signal at 1000 Hz and re-referenced to the electrode placed on the left mastoid, and extracted task trials from −100 to +600 ms with regard to the stimulus onset. We used independent component analysis (ICA)^21^ to remove eye blinking artifacts and movement noises. The grand average event related potentials (ERP) across all the 21 participants are shown separately for the frontal, left parietal, right parietal central, and occipital electrodes (Figure. 4)

**Figure 4.**
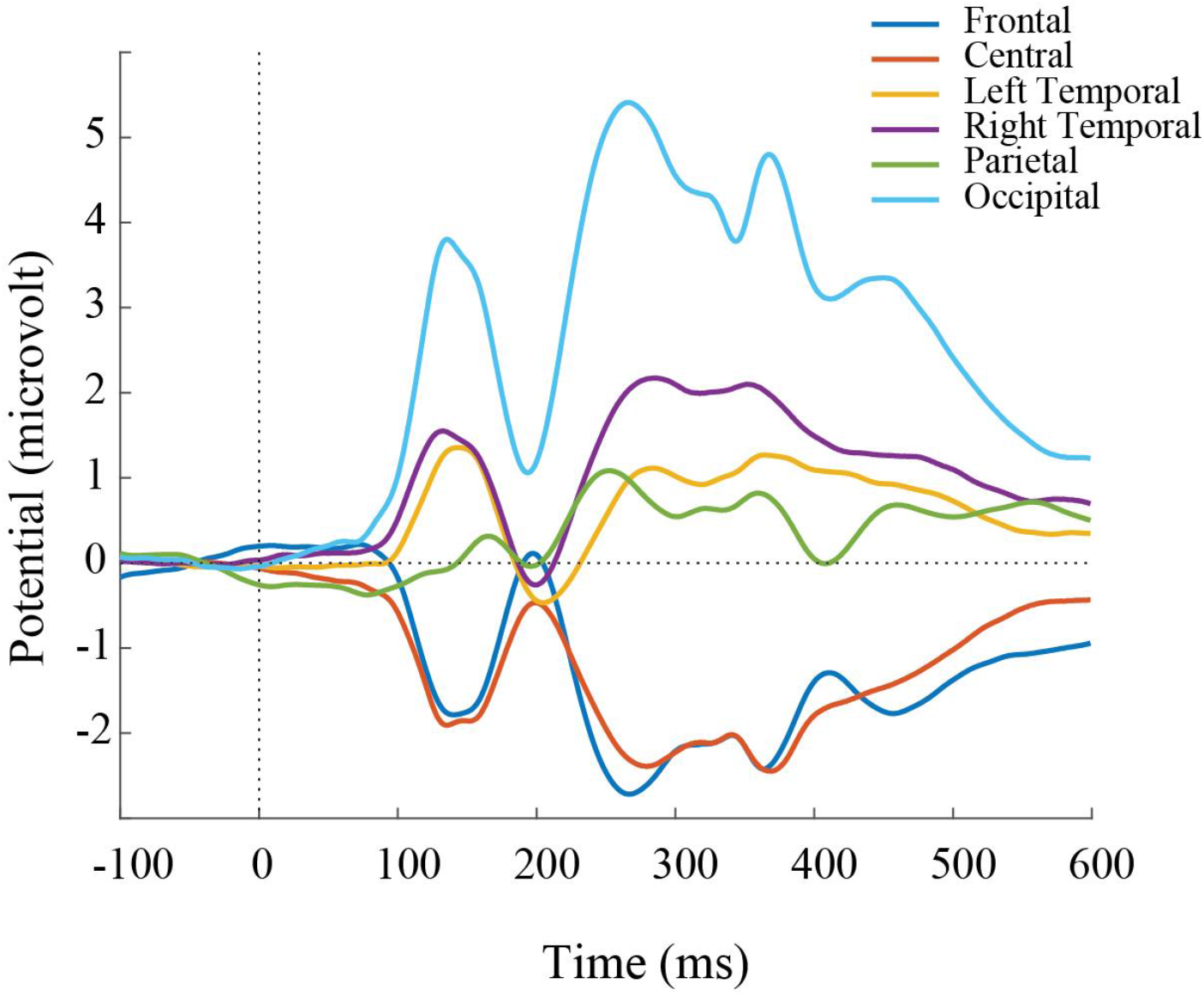
ERPs for 21 subjects at frontal, central, left temporal, right temporal, parietal and occipital electrodes. ERPs for each region are obtained by averaging across all the subjects, all the stimuli, and all the EEG sensors that fall into this region.

#### Multivariate Analysis

To further evaluate the quality of the data, at every timepoint we performed pairwise decoding analysis among the five image categories (one vs. the other four). We used the LIBSVM and RSA-Toolbox^22^ in MATLAB 2016.

There were 800 trials for each subject and category [*800 trials = 5 image identities × 5 variations × 32 repetitions*]. For the pairwise classifications, to compensate for the imbalanced class ratio (i.e. one category vs. four other categories), we selected all the 800 trials from one category and another 800 trials from the other four categories. To Increase signal to noise ratio, we performed sub-averaging on the 800 trials with bin sizes of 5 or 6, leading to 150 pseudo trials being calculated for each category. At each timepoint, we performed leave-one-out cross-validation: for each condition, *N-1* (*N = 150*) pattern vectors were used to train the linear SVM classifier and the *Nth* vector was used for the evaluation. The above procedure was repeated 300 times, every time selecting a different set of random 800 trials.

To assess the significance of classification performance, we used bootstrap resampling of subjects. At each timepoint, we selected samples from subjects with replacement and produced 10,000 bootstrapped samples. The average pairwise classification performance over categories and subjects is shown in Figure. 5 (FDR-corrected over time at P<0.05).

**Figure 5.**
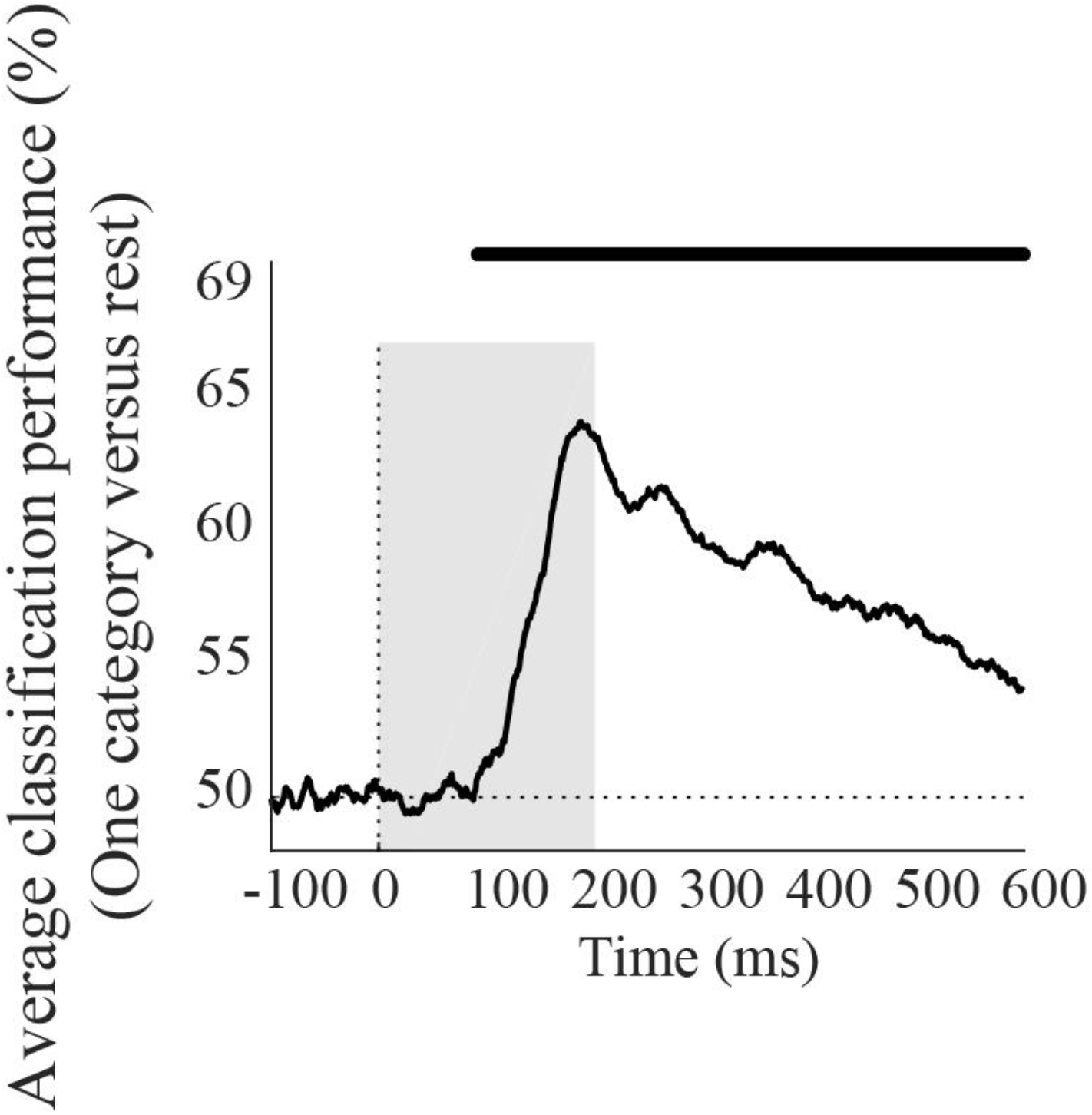
Time course of the average pairwise classification performance showing category selectivity. We performed decoding analysis on each millisecond of EEG data from −100 to +600 ms with respect to stimulus onset. To examine category selectivity, we used a SVM classifier to discriminate between condition-specific EEG sensor patterns. The upper line indicate significant timepoints (Bootstrap resampling test, FDR corrected at 0.05). The gray box indicates the time period of image presentation.

## 5. Code Availability

We used Psychophysics Toolbox ^11–13^ to code the experiment. All EEG preprocessing steps were performed using the EEGLAB toolbox^20^ with MATLAB 2016 ^10^. SPM12^16^ and MATLAB 2016 were used for data preprocessing and applying GLM on the fMRI data. Multivariate pattern analysis of fMRI data was performed using the CosmoMVPA toolbox^18^. The code that was used to convert DICOM files to the *nii.gz* format and generate *.json* files exists in the *code* folder on the repository.

## 6. Acknowledgements

This work was funded by the Iranian National Science Foundation (grant number: 96007585), and was conducted at the National Brain Mapping Laboratory (NBML) in Tehran, Iran.

## 7. Author Contributions

SMK and FE conceived the study and experimental design. FE collected and processed the data. FE analyzed the data with SMK supervision. FE, M.M and SMK wrote the manuscript. M.M helped with the data collection and converted the data to the BIDS format.

